# REINDEER: efficient indexing of *k*-mer presence and abundance in sequencing datasets

**DOI:** 10.1101/2020.03.29.014159

**Authors:** Camille Marchet, Zamin Iqbal, Daniel Gautheret, Mikael Salson, Rayan Chikhi

## Abstract

**Motivation:** In this work we present REINDEER, a novel computational method that performs indexing of sequences and records their abundances across a collection of datasets. To the best of our knowledge, other indexing methods have so far been unable to record abundances efficiently across large datasets.

**Results:** We used REINDEER to index the abundances of sequences within 2,585 human RNA-seq experiments in 45 hours using only 56 GB of RAM. This makes REINDEER the first method able to record abundances at the scale of ~4 billion distinct *k*-mers across 2,585 datasets. REINDEER also supports exact presence/absence queries of *k*-mers. Briefly, REINDEER constructs the compacted de Bruijn graph (DBG) of each dataset, then conceptually merges those DBGs into a single global one. Then, REINDEER constructs and indexes *monotigs*, which in a nutshell are groups of *k*-mers of similar abundances.

**Availability:** https://github.com/kamimrcht/REINDEER

## 1 Introduction

An overwhelming amount of sequencing data is now publicly available in repositories such as the European Nucleotide Archive (ENA) [1] and the Sequence Read Archive (SRA) [2]. They contain a wealth of nucleic sequencing experiments on model and non-model species, tumor biopsies, cell lines, metagenomes. Searching within them, in a holistic setting, would open the tantalizing possibility of making scientific discoveries using world-scale biological data. However, the immense size of these repositories renders any large-scale investigation extremely difficult. SRA hosts around 30 petabases of experiments at the time of writing. Not only is searching within this mass of data impractical, but also merely downloading a copy of ENA or SRA would take in the order of years. Thus, there is a pressing need for solutions that, possibly using a centralized server, would provide search functionality across these databases.

We use the term *dataset* to refer to a set of reads resulting from sequencing an individual sample. Searching for a sequence within a single assembled genome (e.g., with BWA-MEM [3]) or within a single dataset (e.g., with BEETL [4] or SRA-BLAST [5]) is now routine and can be considered to be a well-managed computational problem. Since two decades, searching within collections of assembled genomes (e.g., with BLAST [6] or DIAMOND [7]) is equally practical. However, searching for sequences within collections of (unassembled) datasets is still extremely challenging, and under very active computational investigation [8]. Examples of recent works are HowDeSBT [9], Mantis [10], SeqOthello [11], and BIGSI [12]. They introduced novel *k*-mer (substrings of size *k* in biological sequences) data structures, showing that *k*-mer searches are a useful proxy for exact and approximate sequence search. In a nutshell, state of the art methods are able to determine the presence/absence of any *k*-mer within collections containing up to around 10^4^ indexed datasets. Any longer sequence such as a gene or a transcript can be queried through its constituent *k*-mers. To the best of our knowledge, no method is currently able to record abundances of *k*-mers across collections at this scale.

While nearly all recent works on indexing datasets have focused on recording the presence/absence of *k*-mers, in this work we focus on the more space-expensive matter of indexing their counts, *i.e.*, how many times is a *k*-mer present within each dataset. Foremost, large-scale efforts for indexing abundances in transcriptome collections are currently ongoing. For example the GTEx project [13] hosts a database of transcript-level abundances across 17k tissue samples (as of January 2020). Pan-cancer studies such as TCGA [14] collects the expression of genes, exons, miRNAs, as well as genomic copy-numbers. Those signals are usually recorded at the level of genes or comparatively long genomic regions, yet they would further benefit from nucleotide-level resolution, for example to investigate SNPs and splice junctions. Secondly, metagenomic sequencing is providing a widening window onto the diversity of life on earth. Comparing abundances of contigs, SNPs or species across different metagenomic datasets is a vital step in understanding how community composition varies with context. Thirdly, sequencing now increasingly provides counters or sensors for events: this progression can be seen from CHIP-seq, ATAC-seq, through to single-cell RNA sequencing – after which counts are modelled based on the precise experiment.

We also highlight that merely adapting existing data structures by transforming the 1-bit presence/absence information into a (e.g. 16-bit) counter is unlikely to be a viable strategy. For instance, consider the HowDeSBT data structure [9], a recent technique for indexing the presence/absence of *k*-mers across dataset collections. It saves space by using a single memory location to encode the presence of a *k*-mer across multiple datasets. Yet this scheme cannot be adapted to record abundances, as a *k*-mer may be present in multiple datasets at different abundances, which cannot all be recorded by a single memory location. Likewise, BIGSI [12] uses Bloom filters with 25% false positive rate to encode presence/absence of *k*-mers; extending Bloom filters to support abundance queries (e.g. using Count-Min sketches) at a comparable false positive rate would possibly introduce significant abundance estimation errors.

Here we introduce REINDEER (REad Index for abuNDancE quERy), a novel computational method that performs indexing of *k*-mers and records their counts across a collection of datasets. REINDEER uses a combination of several concepts. The first novelty is to associate *k*-mers to their counts within datasets, instead of only recording the presence/absence of *k*-mers as is nearly universally done in previous works. To achieve this, a second novelty is the introduction of *monotigs*, which allows space-efficient grouping of *k*-mers having similar count profiles across datasets. An additional contribution is a set of techniques to further save space: discretization and compression of counts, on-disk row de-duplication algorithm of the count matrix. As a proof of concept, in this article we apply REINDEER to index a *de facto* benchmark collection of 2,585 human RNA-seq datasets, and provide relevant performance metrics. We further illustrate its utility by showing the results of queries on four oncogenes and three tumor suppressor genes within this collection.

## 2 Problem statement

The questions addressed by REINDEER can be formally framed as follows. Let *S* be a nucleic sequence of arbitrary length (such as a gene, a transcript, or a shorter sequence), and *C* a collection of datasets. The aims are, for each dataset in *C*, to either a) *determine the presence* of *S* or b) *count the number of occurrences* of *S*. In its simplest form, aim a) amounts to returning the subset of datasets in which *S* appears.

In a seminal work, Solomon and Kingsford [15] slightly reformulated aim a) in order to facilitate its resolution. Considering the set *Q* of all *k*-mers extracted from *S*, they propose to instead determine the subset of datasets in which at least *t k*-mers of Q appear, with 0 *<t ≤ |Q*]. This is not strictly equivalent to computing the (exact or approximate) match of *S* within datasets. In this new formulation *k*-mers from *Q* may also be found in different sequences across a dataset, e.g. if ACT and CTG are two *k*-mers seen in two different reads, sequence ACTG is assumed to be present in the dataset regardless on whether it is actually part of a read. Yet this reformulation can be considered as a reasonable approximation. Thus, in this work, we propose a low-footprint index, based on *k*-mer counts, for efficiently supporting a) and b) queries, and more concretely:

1. **Assess***S***’s count in datasets.** Estimate the abundance of *S* in each dataset by reporting the mean count of all *k*-mers from *Q* that are (exactly) present in the dataset.
2. **Assess***S***’s presence in datasets.** Output a list of datasets in which at least *t k*-mers from *Q* appear. Supporting queries of type 1 will be the main improvement proposed by this work compared to the state-of-the-art. In the rest of the manuscript, we will only focus on solving type 1 queries. For queries of type 2 the same principles will apply, as they can be derived from queries of type 1: *S* is considered to be present if and only if it has non-zero abundance.

## 3 Preliminaries

### 3.1 De Bruijn graphs and associated concepts

In this section, we recall the concepts of de Bruijn graphs, unitigs, and compacted de Bruijn graphs. We further describe two recent concepts: spectrum-preserving string sets [16], and the BLight indexing scheme [17].

A *set of reads R* (also called *dataset*) is a set of finite strings over the alphabet {A, C, G, T}. We will consider all the strings of length *k* (called *k-mers*) present in *R*, for some fixed value of *k>*0. For simplicity of presentation, strings are 1-indexed and reverse-complements are omitted from the definitions, but they are taken into account in the canonical way in the software: each *k*-mer is represented by the lexicographically smallest of the forward string and reverse-complemented string.

#### Definition 1. de Bruijn graph (DBG).

*The de Bruijn graph is a directed graph G_k_*(*R*)=(*V,E*) *where V is the set of k-mers that appear in R, and for u,v ∈ V*, (*u,v*)*∈ E if and only if u*[2*,k*]=*v*[1*,k−*1].

Importantly, this definition of the Bruijn graphs is node-centric [18], i.e. the set of edges can be inferred given the nodes. A node-centric de Bruijn graph contains the same information as a *k*-mer set.

We call *unipath* in *G_k_* a maximal-length sequence of distinct nodes *s* =*u*_0_*,…,u_n_* such that, for each 0≤*i≤n*−1, (*u_i_,u_i_*_+1_)∈*E*; in and out-degrees are equal to 1 for each *u_j_* such that 1≤*j≤n*−1; if *n>*1 the out-degree of *u*_0_ is 1; if *n>*1 the in-degree of *u_n_* is 1. Singleton *k*-mers are also unipaths. Intuitively, a unipath is a maximal-length linear portion of the DBG.

#### Definition 2. Unitig.

*The string corresponding to a unipath s* =*u*_0_*,…u_n_ can be defined by concatenating u*_0_ *with all the u_i_*[*k*] *(the k-th letter of u_i_), in order, for* 0*<i≤ n. The resulting string of length k*+*n is called a unitig.*

#### Definition 3. Compacted de Bruijn graph.

*The directed graph where nodes are unitigs, and edges correspond to* (*k−*1)*-overlaps between two nodes sequences, is called a compacted de Bruijn graph.*

Contrary to a regular DBG, a compacted DBG has nodes of various lengths (≥*k*). In the following, unless explicitly mentioned, the objects referred to as “de Bruijn graphs” are compacted de Bruijn graphs. Even though compacted DBGs are technically not DBGs, they do equivalently represent the same *k*-mer set. Each *k*-mer has a count corresponding to the number of times it appears in the reads of a dataset, which can be obtained using a *k*-mer counter [19].

#### Definition 4. Unitig abundance.

*We call the unitig abundance c*(*u*):

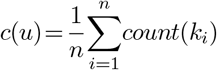

*for k*_1_*,…,k_n_ the k-mers of the unitig u of length n*+*k−*1.

#### Definition 5. Union De Bruijn graph.

*Given a collections of de Bruijn graphs* 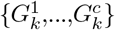, *we call* union de Bruijn *graph the de Bruijn graph which has a node set equal to the union of the nodes set of* 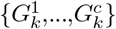*, and edges correspond to* (*k−*1)*-overlaps between k-mers of the node set.*

It follows that a union DBG also represents a *k*-mer set (the union of *c k*-mer sets). We now recall the recent concept of spectrum-preserving string sets [16], in order to draw new bridges between previous works, and also with our contribution.

#### Definition 6. Spectrum-preserving string set (SPSS) [16].

*Given a multiset of k-mers X, a set of strings S of length ≥ k is a SPSS representation of X if the multiset of all k-mers from S is exactly X.*

In particular, a SPSS can be used to represent the node set of a DBG. In such case, the SPSS represents a set (i.e. a multiset with no duplicates) as all *k*-mers are distinct. Unless explicitly noted, we will further consider in the rest of the article that all SPSSs represent sets and not multisets.

Given a set of *k*-mers *X*, the trivial set of all *k*-mer strings of *X* is itself a SPSS of *X*, albeit not a very space-efficient one. The set of unitigs constructed from *X* is also of SPSS of *X*. We will discuss more types of SPSSs below.

### 3.2 Spectrum-preserving string sets in relation to k-mer indexing

One of the fastest and most space-efficient techniques available to index and associate information to *k*-mers is BLight [17], a hash table scheme tailored to large sets of *k*-mers. Given a collection of datasets, the input data given to BLight is a SPSS of the *k*-mers from the union de Bruijn graph of this collection. In its inner steps, BLight relies on a particular type of SPSS, where in each sequence of the SPSS all *k*-mers need to have the same minimizer.

#### Definition 7. Minimizer [20].

*The m-minimizer of a sequence S is the smallest substring of size m in S, according to some ordering.*

Here *m*-minimizers are computed on *k*-mers, hence *m<k*, and the order on *m*-mers is given by a hash function (i.e., not the lexicographic order).

#### Definition 8. BLight indexing scheme [17].

*Given a set of k-mers X, let S be a SPSS of X such that in each string s S, all k-mers in s have the same m-minimizer. Then BLight uses S to compute a static, collision-free hash table where the set of keys is X.*

In the original BLight article, the SPSS of *X* is the set of all the super-*k*-mers found in the unitigs, as defined below.

#### Definition 9. Super-*k*-mer [19].

*Given a sequence x, a super-k-mer is a substring of x of maximal length such that all k-mers within that substring have the same m-minimizer sequence.*

#### Observation 1.

*Given a set of k-mers X, the set of all super-k-mers constructed from the unitigs of the DBG of X is a SPSS of X.*

Concretely, the original version of BLight takes as input the unitigs of the DBG of *X*, and transforms them into another SPSS: the super-*k*-mers of unitigs, partitioned according to their minimizers. However, it is important to note that this is not the only possible SPSS scheme that can be used by BLight. In fact, in this article we will use another one.

In the inner workings of BLight, querying a *k*-mer *x* consists in first computing the minimizer of *x* in order to identify in which partition it is expected to be found. Then a minimal perfect hash function [21] associates *x* to its position within a sequence of the SPSS. The position is then converted into a number that indicates a location in memory, that can be used to associate some piece of information to each *k*-mer. Of note, BLight only records information about the set of *k*-mers itself, but has no additional information such as their presence/absence within a collection of datasets, nor their counts. Another data structure will be needed along with BLight to record such information, which is in fact the purpose of REINDEER.

So far, we have reviewed the following SPSSs for a set of *k*-mers *X*: *X* itself, the unitigs of *X*, any set of super-*k*-mers that together contains all *k*-mers of *X* (such as the super-*k*-mers of the sequencing reads where *X* originated from), and the super-*k*-mers of the unitigs of *X*. To this list we can add the recently-introduced (and equivalent) concepts of UST and simplitigs [16, 22]. They are SPSSs that aim to minimize their total number of nucleotides. Note that contigs resulting from a genome or transcriptome assembly of *X*, unlike unitigs, are not a SPSS as they typically discard some of the *k*-mers to generate consensus sequences. Table 1 recapitulates all SPSS schemes (monotigs will be defined later in the article). In the next section, we will show that the choice of SPSS is important for our specific application. We will introduce a new SPSS that is better suited to the purpose of indexing collection of datasets.

**Table 1:**
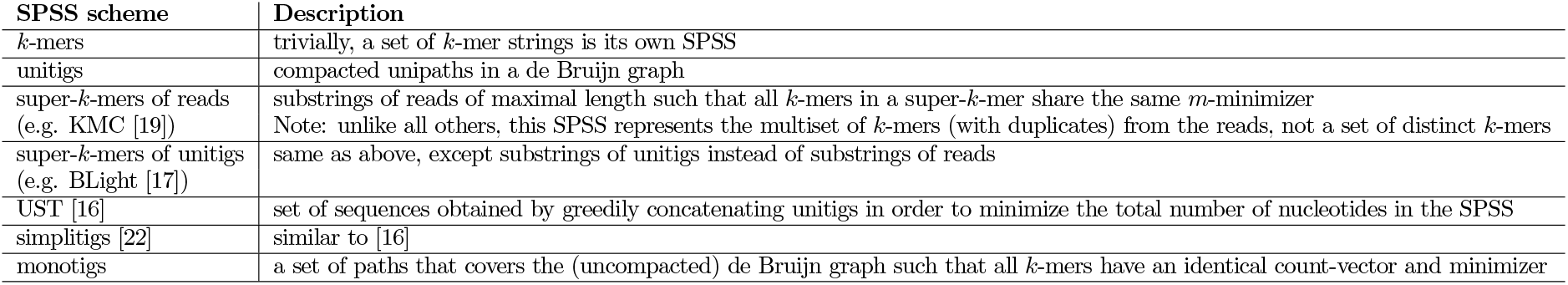
Categories of spectrum-preserving string set schemes known from previous literature (and monotigs, introduced in this article). See also Fig. 1 for an example of each category.

| SPSS scheme | Description |
| --- | --- |
| $k$-mers | trivially, a set of $k$-mer strings is its own SPSS |
| unitigs | compacted unipaths in a de Bruijn graph |
| super-$k$-mers of reads<br>(e.g. KMC [19]) | substrings of reads of maximal length such that all $k$-mers in a super-$k$-mer share the same $m$-minimizer<br>Note: unlike all others, this SPSS represents the multiset of $k$-mers (with duplicates) from the reads, not a set of distinct $k$-mers |
| super-$k$-mers of unitigs<br>(e.g. BLight [17]) | same as above, except substrings of unitigs instead of substrings of reads |
| UST [16] | set of sequences obtained by greedily concatenating unitigs in order to minimize the total number of nucleotides in the SPSS |
| simplitigs [22] | similar to [16] |
| monotigs | a set of paths that covers the (uncompacted) de Bruijn graph such that all $k$-mers have an identical count-vector and minimizer |

## 4 SPSSs for indexing *k*-mer counts

Due to the stochasticity of sequencing coverage, genomes and transcriptomes are not evenly covered by sequencing reads. Thus, *k*-mers coming from regions that have the same copy-number (in genomes, – or abundance, in transcriptomes) often have close but non-identical counts. Yet, if those *k*-mers had identical counts, such redundancy could be exploited to save space when indexing counts. While it would be unrealistic to assume that *k*-mer counts are constant across long genomic regions of same copy-numbers (or of same abundances across a transcript), we will assume it for shorter regions (unitigs) throughout the rest of the article.

### Assumption 1.

*Within a dataset, the counts of k-mers that are part of the same unitig are assumed to all be identical.*

We argue that Assumption 1 yields a robust approximation of *k*-mer counts. To support this claim, we computed the Pearson (*r*) and Spearman (*ρ*) correlation coefficients of true *k*-mer counts versus counts averaged per unitig, across 15 RNA-seq datasets, resulting in *r* =0.999 and *ρ* =0.910 on average (see Figure S2 in the Appendix).

### 4.1 Count vectors and SPSSs

Using the indexing scheme presented in the previous section (a SPSS indexed by BLight), our aim is to associate to each *k*-mer its abundances in datasets.

#### Definition 10. Count-vector.

*Given an ordered list of datasets and a k-mer x, the* count-vector *of x is a vector in which the integer at position i represents the number of times x is seen in the i-th dataset.*

Our initial motivation is to seek a SPSS scheme where a single count-vector can be associated to each string of the SPSS. In other words, for each string *s* of the sought SPSS, all *k*-mers in *s* need to have the same count-vector.

#### Observation 2.

*Under Assumption 1, given a super-k-mer constructed from the unitigs of a union DBG, it is possible that not all k-mers in the super-k-mer have the same count-vector.*

We show a proof of Observation 2 in Appendix, illustrated by Figure 2. It is immediate that a possible instance of the sought SPSS is *X* itself. Since each string of that SPSS only contains a single *k*-mer, a single count-vector is associated to each element of the SPSS. However this is not a scalable solution, as thousands of mammalian-sized datasets contain billions of distinct *k*-mers, leading to prohibitively high memory consumption.

Yet recall that BLight allows to index any SPSS, as long as it satisfies the minimizer condition of Definition 8. It turns out, perhaps surprisingly, that none of the SPSS schemes of the previous section would be suitable for associating elements to single count-vectors. We describe why below.

Most of the time, the *k*-mers within a unitig do not share the same minimizer, thus unitigs cannot be used as a SPSS for BLight. This is why in the original BLight scheme, super-*k*-mers are extracted from the unitigs in order to construct a suitable SPSS. Even so, a second issue occurs: in the union DBG, two *k*-mers coming from two different datasets can overlap over *k* 1 nucleotides, forming in a unitig a (*k* +1)-mer that exists in no single dataset. We will call such unitigs *chimeric* (see Figure 2, and leftmost unitig in Figure 1). It is clear that *k*-mers within chimeric unitigs do not all have identical count-vectors. This leads to the impossibility of associating a single count-vector to chimeric unitigs or even to super-*k*-mers of chimeric unitigs.

**Figure 1:**
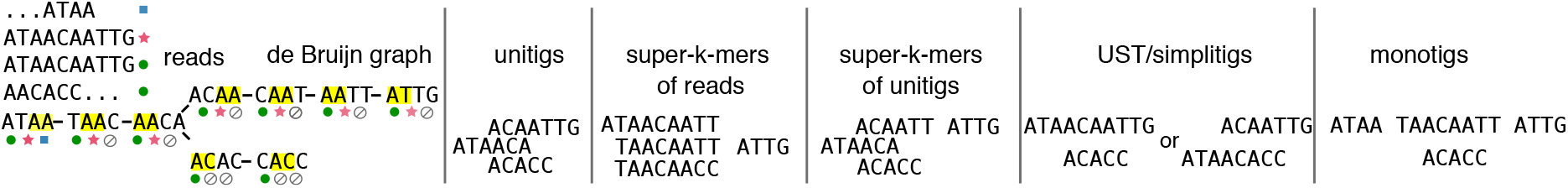
Example of a de Bruijn graph (left part) and various SPSS schemes over the set of nodes of the graph (rightmost panel). There are three datasets (circle/star/square symbols). In this first panel, the minimizer of each *k*-mer is highlighted in yellow (*k* =4, *m* =2). Dots below *k*-mers symbolize their non-zero abundance (filled symbols) or absence (Ø) across the three datasets. In the five other panels, sequences within each SPSS scheme are roughly represented according to the positions of constituent *k*-mers in the DBG, only for visual indication. Observe that monotigs are the only SPSS scheme where each sequence can be associated to a single combination of colored dots.

**Figure 2:**
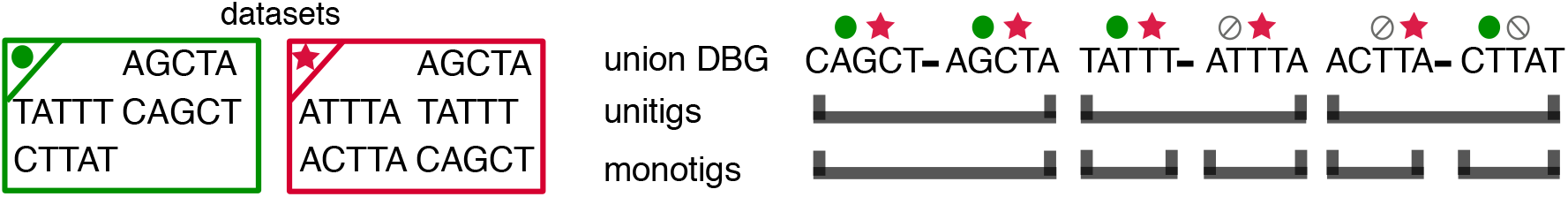
Left: 2 datasets and their *k*-mers (*k*=5). Right: a union DBG is built over these *k*-mers, and contains 3 unitigs (top row). Symbols (e.g. circle, star) above each *k*-mer symbolize the per-dataset abundances. In the left-most unitig, the two *k*-mers have the same abundance information. Thus the monotig is here equivalent to the unitig. In the center and right-most unitigs, *k*-mers that have contrasted abundances share a *k* 1 overlap and can be compacted. On the contrary, monotigs will stop at differences between the coverage profiles of overlapping *k*-mers.

Of note, the other SPSS schemes UST and simplitigs aim to merge unitigs as much as possible. These schemes suffer from the same issue of having multiple count-vectors within a single sequence, at an even more extreme level than with single unitigs.

### 4.2 Monotigs

As we saw in the previous section, super-*k*-mers constructed from the unitigs of the union DBG were likely to be a suitable SPSS, except that Observation 2 shows that a single count-vector cannot always be associated to each of them. However, this issue does not arise when considering a single dataset at once. This allows us to introduce the new concept of monotig.

#### Definition 11. Monotig.

*A* monotig *is the sequence of a path in the union DBG in which all constituent k-mers have the same count-vector and the same minimizer.*

Observe that monotigs are not necessarily substrings of super-*k*-mers of unitigs as they can in principle span multiple unitigs (see e.g. Figure 3 step 2/). The set of monotigs forms a partition of the nodes of the DBG. Note that contrary to other SPSSs, due to the count vector constraint, monotigs are not solely defined using *k*-mers of the de Bruijn graph.

**Figure 3:**
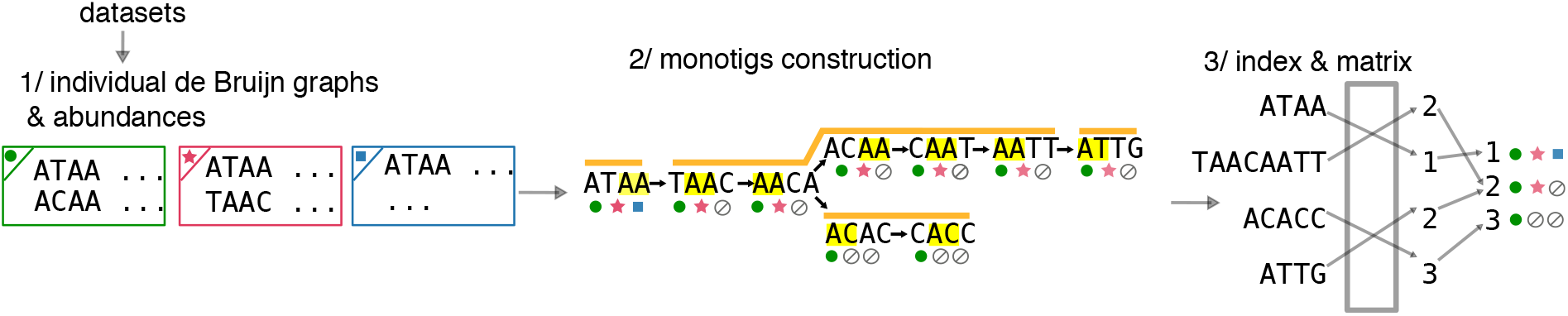
Overview of REINDEER index construction. At step 1/, for each dataset, the (compacted) de Bruijn graph is given as input along with the count of each unitig. At step 2/, monotigs (corresponding to paths in orange) are computed on the *k*-mers of the union De Bruijn graph of all datasets, and these monotigs are indexed. During step 3/ each monotig is associated (through an integer array) to a row in the de-duplicated count-vector matrix (rightmost matrix). In this matrix no two rows are equal, and in practice each row is compressed.

## 5 REINDEER data structure

Given a collection of datasets, REINDEER is an index that associates to any *k*-mer its counts in all datasets. As a byproduct of recording counts, it is also possible to query the presence/absence of *k*-mers within these datasets. This makes REINDEER a memory-efficient representation of colored de Bruijn graphs, such as other recent methods in the literature [10, 23, 11], and in particular Bifrost [24] which indexes unitigs through the minimizers of their constituent *k*-mers. But note that REINDEER does more than representing colors (it also records unitig-averaged abundances), and it is static (unlike Bifrost) and not particularly designed for efficient graph traversal.

In line with recent works [10, 11, 9], REINDEER conceptually transforms an input collection *C* of datasets into a set of multisets of *k*-mers. As a pre-requisite, a compacted DBG needs to be built for each dataset (e.g. using BCALM2 [25]). REINDEER then takes as input the collection of compacted DBGs, and a single count value per unitig present in each DBG. It represents implicitly the union DBG of all the compacted DBGs, which is never explicitly constructed in memory. To summarize, REINDEER is built in three steps, illustrated in Figure 3:

1. For each dataset, build an individual DBG that also records *k*-mer counts, averaged per unitig.
2. Construct monotigs (according to the greedy algorithm of Section 5.1) to index all *k*-mers from the union DBG of all datasets.
3. Associate count-vectors to monotigs, and further discard duplicated count-vectors. The following three sections describe monotig construction, and the association of monotigs to count-vectors in more details.

### 5.1 Construction of the monotigs

We propose a greedy algorithm to compute a set of monotigs efficiently. The algorithm takes as input a set of compacted DBGs (one DBG per dataset). First, super-*k*-mers of unitigs are computed for each dataset. The rationale for using super-*k*-mers instead of *k*-mers is to save space, as opposed to having to store all *k*-mers on disk. Each super-*k*-mer records its count in the DBG it originates from (there is a unique count, according to Definition 4). The super-*k*-mers are partitioned into files according to their minimizers. At this step, the same *k*-mer may be present in multiple super-*k*-mers within a file (in fact, the *k*-mer is repeated as many times as the number of datasets where it is present). Then, for each minimizer file, super-*k*-mers are broken into their constituent *k*-mers. The *k*-mers and their count-vectors are inserted into a hash table^1^. When a *k*-mer is first inserted, it has a count-vector that records the abundance of its originating super-*k*-mer in the corresponding dataset. When a duplicate of a *k*-mer already in the hash table is met by parsing another super-*k*-mer (i.e., the same *k*-mer coming from another dataset), the associated count-vector is updated by adding the abundance of the other dataset.

In each minimizer file, monotigs are constructed by greedily merging *k*-mers having identical count-vectors and a (*k* − 1) overlap. Concretely, each *k*-mer at an extremity of a monotig queries the hash table for the existence of a (uncompacted) DBG neighbor (e.g. for a right extremity, neighbors are the (*k*−1)-suffix of the *k*-mer appended with either A, C, T, or G). A monotig is extended by an arbitrary neighbor if one is found. The procedure is repeated until no more extension is possible. Observe that each minimizer file can be processed independently and in parallel.

### 5.2 Construction of the count-vector matrix

Once all monotigs are computed, they are given as input to BLight. The BLight scheme is slightly modified in order to associate a single memory location to all *k*-mers within a monotig, instead of one memory location per *k*-mer. This is done in a straightforward way by returning the monotig’s identifier instead of the position of the queried *k*-mer within that monotig.

Once the BLight index is built, a *count-vector matrix M* is created such that its rows are the monotigs count-vectors, organized according to the order of the monotigs referenced by the values of the BLight hash table. In other words, the value *M*[*i*][*j*] in this matrix gives the count of all *k*-mers within the *i*th monotig in the *j*th dataset. The matrix is stored in a row-major order; each row can be accessed efficiently during queries.

### 5.3 De-duplication of rows in the count-vector matrix

Colored de Bruijn graphs and similar works initially used matrices to record colors (i.e., presence/absence information) of *k*-mers within a de Bruijn graph [23]. More recently, color equivalence classes have been introduced to decrease the redundancy in the matrix [26, 10]. We apply a similar idea to save space, by de-duplicating rows of the count-vector matrix.

Simply put, we do not store the count-vector matrix, but instead we store a transformed version (the *de-duplicated count matrix*) which contains only distinct rows. In addition, we maintain an integer vector that associates to each monotig its corresponding row in the de-duplicated count-matrix.

### 5.4 Query

Queries in REINDEER are performed as follows. A query sequence *S* of size *L* is decomposed into its constituent *k*-mers, which are then individually queried in the index. Behind the scenes, the index converts each *k*-mer into a monotig identifier, used to access the corresponding row of the de-duplicated count-vector matrix, then retrieves the counts associated to this *k*-mer in all datasets.

Note that the presence/absence of a query *k*-mer *x* can be directly inferred from the count-vector matrix. Indeed, if *x* has a non-zero count in dataset *j*, it means that *x* is present in one of the unitigs of the DBG of dataset *j* and thus in some monotig *i* of the union DBG. Thus, the presence of *x* can be deduced by computing whether *M*[*i*][*j*] *>*0.

To determine the abundance of *S* in each dataset, we first check whether *S* is *present* in the dataset: whether at least *θ* of the constituent *k*-mers of *S* have non-zero abundance in the dataset (by default, *θ* =40%). If REINDEER is asked to only determine the presence/absence of *S*, the query stops here. For abundance queries, a matrix of integers *Count* is recorded, of size *m × c* where *m* =*L − k*+1 is the number of *k*-mers in *S* and *c* the number of datasets. Each value *Count*[*i*][*j*] corresponds to the count of the *i*-th *k*-mer of *S* inside dataset *j*.

Instead of directly reporting the matrix *Count*, we perform an additional step that de-duplicates identical values within each column. For example if in a given dataset *j* all the counts of *k*-mers present in *S* are identical (|{*Count*[*i*][*j*]*,ᗄi* s.t. *Count*[*i*][*j*] *>*0 =1), then we report a single count value for this dataset. On the contrary, if not all *Count*[*i*][*j*]s are identical at a fixed *j*, then a list containing all seen count values is reported for dataset *j*. This second case happens when *k*-mers from *S* fall into different unitigs of dataset *j*, and these unitigs have different abundances (as defined by Definition 4). Thus the final query result for a given sequence *S* is a tuple containing *c* (possibly single-valued) lists.

An example of REINDEER output is given in Figure S1 in Appendix. Relying on BLight allows for fast queries and, unlike other methods based on the Burrows-Wheeler transform, a constant (and small) number of memory accesses are performed. Query complexity for the presence/absence (or abundance) of a sequence of length *L* (*L> k*) in an index of *c* datasets is in time (*L×c*). Although queries that scale linearly with the number of datasets are not ideal theoretically, in practice engineering mitigates this problem. Indeed, grouping all datasets in a single index enhances cache-locality and allows rapid queries by reaching all datasets for a given *k*-mer in less than *c* RAM accesses. Other data structures (Mantis [10], Bigsi [12]) also have this query complexity. Moreover, a main focus of REINDEER is to have a low-footprint during queries. This is possible since REINDEER benefits from compression, which allows loading in-memory all datasets information at once while reducing the memory footprint.

### 5.5 Implementation details

#### 5.5.1 REINDEER index

When loaded in memory, the BLight index is not further compressed and the abundance matrix is compressed with RLE. Both are compressed (with `gzip`) when dumped to disk, and are reloaded into memory (de-gzipped) when doing a query.

#### 5.5.2 Discretization of counts

If the accuracy of *k*-mer counts is not critical, then encoding counts using reduced integer precision leads to more effective de-duplication of the count-vector matrix. In REINDEER, we implement two different techniques. 1) Applying a function (termed *discretization*) that encodes counts from 0 to 50,000 in 8-bit integers, ensuring that discretized counts are at most 5% different from the real raw counts^2^. 2) A log_2_ transformation of counts (formally, max(0, 『log_2_(*count*)』)) yields an even more extreme discretization (in terms of loss of precision). An example of these transformations is shown in Table 2. We emphasize that these are not the only possible discretization schemes. Further investigations would also be necessary to determine their impact on biological queries. For example, a 5% difference in large counts (e.g. 10000 vs 10500) may turn out significant in differential expression analysis, but would be masked by discretization.

**Table 2:** Types of count-vectors associated to *k*-mers.

| Type | Sample values for 4 datasets |  |  |  |
| --- | --- | --- | --- | --- |
| Raw counts | 27 | 0 | 101 | 40812 |
| Discretized counts | 27 | 0 | 102 | 41000 |
| $\log_2$-transformed counts | 4 | 0 | 7 | 16 |
| Presence/absence | 1 | 0 | 1 | 1 |

#### 5.5.3 Low-memory de-duplication of rows in the matrix

We iteratively compute the count-vectors of monotigs and compress them with run-length encoding (RLE)^3^, and then partition the whole multi-set of count-vectors, using hashing, into files stored on disk. When a count-vector is written to disk, its monotig identifier given by BLight is also recorded next to it. Then, reading each file separately, we select a single representative of each vector by inserting it into an efficient dynamic hash table^1^. Once a partition is processed, we write the set of de-duplicated count vectors to disk, and record the mapping between monotig indices and their positions in the de-duplicated count-vector matrix.

#### 5.5.4 Query speed-up and low-memory footprint

In order for a query length to not increase the memory footprint, we only decompress and read a single count-vector at a time, i.e. we process *k*-mers sequentially. We also take advantage of the fact that consecutive *k*-mers *x,y* in a query are also likely to be present in the same monotig. If it is the case, we directly copy the information of *x* for *y* instead of again interrogating the de-duplicated count-vector matrix.

## 6 Results

In their work, Solomon and Kingsford [15] indexed 2,652 human RNA-Seq samples to demonstrate the scalability of their approach. Since then, other competing methods have also used this dataset. It was further reduced to 2,585 datasets to remove experiments containing short reads that could not be indexed with 21-mers [9], and has become a *de facto* benchmark. We use results from a recent and exhaustive review [8] in order to compare REINDEER with the state-of-the-art in indexing collections of datasets. Despite not having run all methods on the same machine, the comparison of *space* usages for each data structure will nevertheless be accurate; results regarding execution times will certainly be machine-dependent but orders of magnitudes are expected to be roughly preserved.

We compare our approach with 5 others: Solomon and Kingsford’s original Sequence Bloom Tree (SBT) [15]; HowDeSBT, the most recent improvement of SBT [9]; other hashing-based methods (BIGSI [12], SeqOthello [11]); and a recent method relying on a de Bruijn graph (Mantis [10]). Among those approaches, only REINDEER and Mantis support exact *k*-mer queries, all the others are probabilistic, meaning that a given *k*-mer may be said to be present, while it is not. REINDEER is the only index that provides count information (as opposed to just presence/absence). We provide scripts and guidelines^4^ to reproduce the following results.

### 6.1 Index construction

#### 6.1.1 Comparison to state-of-the-art

We first build REINDEER indexes on the 2,585 datasets and filter out the rare *k*-mers on the same criteria as the other methods^5^, resulting in a total of ~ 3,755 millions of distinct *k*-mers to index after filtering. We then compare our performance metrics to those reported by other tools on the same dataset [8]. Several construction modes are assessed: indexing presence/absence information only (for the sake of comparison only as our contribution consists in providing counts); and indexing counts in different modes (normal, discretized, log_2_). In order to be comparable to other results, we used a *k* value of 21 (and *m* = 10). Other parameters in REINDEER are set to default values, with commit version d8a9eec.

Table 3 shows the resources required at construction. Globally, REINDEER requires comparable resources as the existing methods, yet they only support presence/absence queries. REINDEER requires more disk space and the index is about as space consuming as Mantis, which also offers exact *k*-mer queries. However REINDEER was not conceived with presence/absence queries in mind, but for abundance queries. In such a case we see that the index is very efficient as it is about 2 to 4 times more space-expensive than presence/absence methods (yet with many of them having false positives). REINDEER space consumption can be additionally reduced by a third when storing approximate counts.

**Table 3:** Index construction resource requirements on 2,585 RNA-seq datasets on REINDEER and as reported in the literature for the other indexes. The “Max Ext. Memory” column reports the maximal external (i.e., disk) space taken by the method. Execution times of methods are reported in wallclock hours. “Index size” indicates the final REINDEER index size as stored on disk (i.e., the sum of three components: the index of monotigs, the abundance matrix and the vector mapping monotigs to matrix rows). N/A indicates that the value was not reported in the article of the given method.

| Tool | Max Ext. Memory (GB) | Time (h) | Peak RAM (GB) | Index Size (GB) | Counts (Y/N) |
| --- | --- | --- | --- | --- | --- |
| SBT | 300 | 55 | 25 | 200 | N |
| HowDeSBT | 30 | 10 | N/A | 15 | N |
| Mantis | 3,500 | 20 | N/A | 30 | N |
| SeqOthello | 190 | 2 | 15 | 20 | N |
| BIGSI | N/A | N/A | N/A | 145 | N |
| REINDEER - presence/absence | 6,800 | 40 | 27 | 36 | N |
| REINDEER - raw counts | 7,000 | 45 | 56 | 52 | Y |
| REINDEER - discretized | 7,000 | 40 | 44 | 44 | Y |
| REINDEER - $\log_2$ | 7,000 | 64 | 32 | 34 | Y |

We also built REINDEER index on a different data type (metagenomic samples), and show the results in Table S3 in Appendix.

#### 6.1.2 Scalability and influence of *k*

In this paragraph we assess the performance of constructing REINDEER on several datasets sizes and *k* sizes. In order to fairly compare the different runs, we selected a set of 2,512 datasets that have reads of lengths ≥31 bases among the 2,585 previously mentioned datasets. We assessed the scalability of the REINDEER index (for raw counts) on 10 to 2,512 datasets (Table 4). The resources required grow roughly linearly with the number of distinct *k*-mers. The performance of REINDEER can be impacted by the chosen *k*-mer size. Intuitively, longer *k*-mers can lead to larger monotigs, thus to a reduction of REINDEER index size. We show the behavior of REINDEER on several *k* values in Appendix Table S1. Monotigs are longer with *k* = 31 (35 bases on average, compared to 22 bases on average for *k* = 21). Additional metrics (number of monotigs, de-duplication and RLE gains) can be found in Table S2 in the Appendix.

**Table 4:**
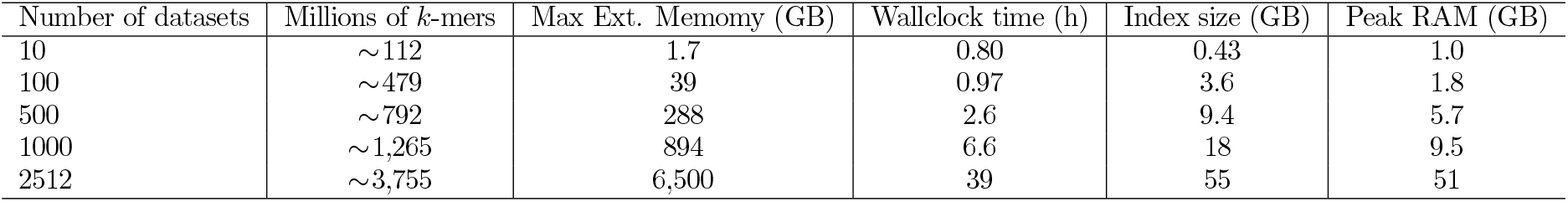
Performance of REINDEER (raw counts, *k* =21) on subsets of datasets from the 2,512 human RNA-seq datasets. Columns are: number of distinct *k*-mers, maximum space usage on the disk, construction wall-clock time (20 threads), index size after construction, and memory peak during construction.

#### 6.1.3 Comparison with the Jellyfish and other hash tables

We selected Jellyfish [27] for a comparison with REINDEER because it is one of the most straightforward existing alternatives: Jellyfish implements a hash table that associates to each *k*-mer its count within a dataset. We compare REINDEER and Jellyfish by estimating the space consumption of Jellyfish per key to index, and let aside the abundance storage. The space usage of recording 4 billion distinct *k*-mers with Jellyfish is approximately 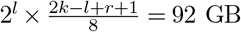 (according to Jellyfish manual, with *l* = *log*_2_(*K/*0.8), *r* = [*log*_2_(62)]−1, *k* = 21 and *K* = 4.10^9^). In comparison, REINDEER indexed monotigs and their associations to memory locations using roughly 15 GB of memory, that is, a 84% space saving relative to Jellyfish.

We also briefly compare the monotigs indexing part of REINDEER with two other hashing schemes: Short Read Connector (SRC) [28] and the original BLight scheme [17]. SRC indexes *k*-mers directly, and was shown to outperform classical hashing techniques (such as C++ unordered maps). As described above, BLight indexes super-*k*-mers of unitigs. We experimentally tested the original BLight implementation on 4 billion distinct *k*-mers, resulting in 15 GB of memory usage, similarly to REINDEER. This demonstrates that switching the SPSS strategy from unitig super-*k*-mers to monotigs has no negative impact on the index size. On the other hand, SRC used 23 GB of memory.

### 6.2 Query performance

We then estimated the abundance of some test sequences in the 2,585 datasets using the REINDEER index. Queried sequences were randomly sampled from the RefSeq human transcripts. Batches of 10, 100, 1000 and 10,000 sequences were created in order to study query scalability. The mean sequence size over the batches was 3,341 nucleotides. Each batch type was repeated 10 times with random sampling, and results show the mean, max and min query wallclock times for each batch type (using 20 threads). Results are shown in Table 5. The query wallclock time can be divided into a first step in which REINDEER loads the index into memory, and a second step in which *k*-mers from each query are queried in the REINDEER structure. Final results are computed and written to an output file. We used the same parameters as for the index construction (*k* =21*,m* =10, −S=75%, with −S being the minimum percentage of found *k*-mers required to return a hit in a dataset. All the experiments were performed after first warming the cache with the index. The peak of RAM consumption is also represented in Table 5, and fluctuates only moderately as it represents the size of the un-gzipped index in memory, plus an additional structure corresponding to the *n* count-vectors of size 2585 for the *n k*-mers of a query sequence.

**Table 5:** Performance of REINDEER in terms of index loading time, query time, and peak memory during query, for different human transcripts batch sizes.

| Batch size | Index loading (s, wallclock)<br>mean/min/max | Sequences query and I/O (s, wallclock)<br>mean/min/max | Peak RAM (GB) |
| --- | --- | --- | --- |
| 10 sequences | **867**/836/916 | **92**/89/95 | 84.8 |
| 100 sequences |  | **149**/139/160 | 88.4 |
| 1,000 sequences |  | **394**/362/410 | 83.8 |
| 10,000 sequences |  | **1042**/986/1095 | 87.0 |

As measurements of query times are architecture-dependent, comparisons with other methods are given here as a rough estimate. According to a recent benchmark [29], SeqOthello and Mantis performed queries on 1000 sequences in a similar experiment in around a minute. For the same task, SBT would take an hour or so, and the more recent HowDeSBT around 5 minutes.

### 6.3 Abundance matters: an example with oncogenes

Although individual cancers have highly variable origins, genes from the oncogene and tumor suppressor families often show abnormal expression in cancer tissues. Certain oncogenes have low expression in healthy tissues but are over-expressed in cancer, while certain tumor suppressors vary in the opposite fashion.

Some oncogenes are normally expressed in healthy patients but can be over-expressed in cancer patients. With our REINDEER index we will demonstrate a simple *in silico* experiment involving theses abundances.

For genomic data, where reads are used to reconstruct the genome, *k*-mer-indexes can be used to query presence of alleles, but (modulo copy-number differences) all of them should be there at the genome-wide average abundance. In stark contrast, the situation is different for RNA-seq. There, every transcript may be expressed at a different abundance, and one might look for different patterns of expression. Given an index of an expression archive, it becomes possible to perform hitherto impossible queries. One can take an array of genes/transcripts, and collect an abundance signature across thousands of datasets, and then perform unsupervised clustering to look for patterns. One can pick a transcript of interest (e.g., from an oncogene) and compare the abundance of that and a control transcript, across all datasets, or maybe compare this query across tumour and normal samples (assuming appropriate metadata is available).

As a proof of concept we selected four oncogenes well known for their role in breast cancer (ERBB2, FOXM, MYC, PIK3CA), as well as three tumor suppressor genes (BRCA1, PTEN, TP53) [30, 31]. We launched REINDEER on the longest transcript of those genes. The transcript was split in 100bp long sequences, to be able to take into account matches on a few exons. We required at least 78% of the *k*-mers to be found in the split sequences. We averaged the counts returned by REINDEER in each split sequence and kept the maximal value among them. The distribution in each dataset, for which counts are reported, are displayed in Figure 4. We recall that our protocol is only a demonstration of what could be done with REINDEER. Obtaining robust quantifications, in a similar way as e.g. Kallisto [32] but on thousands of datasets in a single query, is an important direction left for future work.

**Figure 4:**
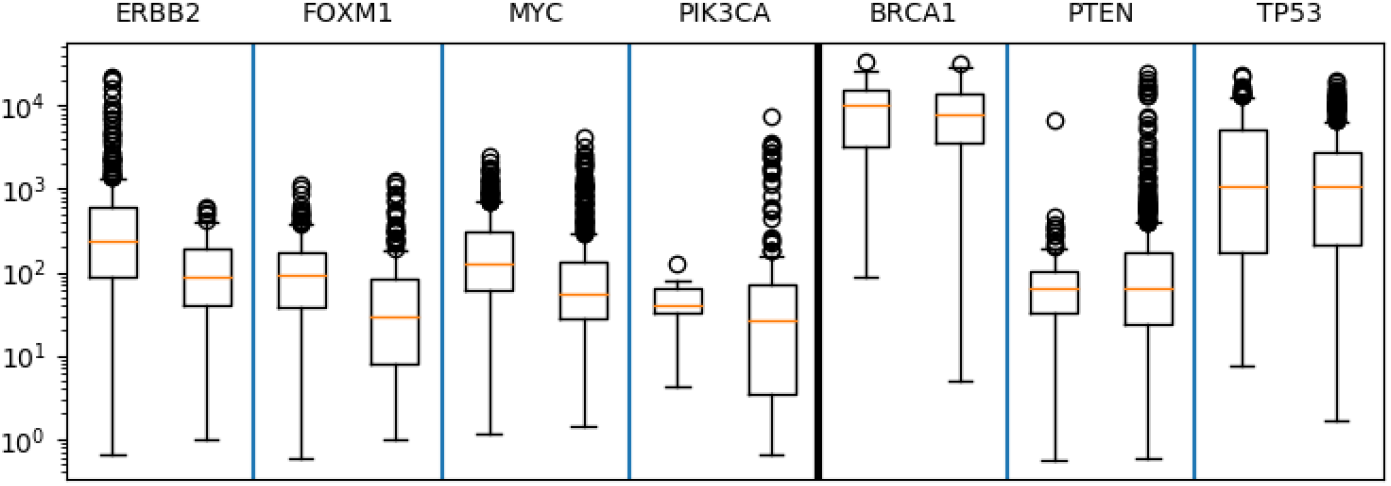
Distribution of the average counts reported by REINDEER in the longest transcript, both in “cancer-related” datasets (left barplot in each panel) and in “non cancer-related” (right barplot) datasets for four oncogenes (ERBB2, FOXM1, MYC, PIK3CA) and three tumor suppressor genes (BRCA1, PTEN, TP53) considered. The median fold changes are respectively 2.1, 2.4, 1.9, 1.8, 1.4, 1.1, 0.93.

Even with crude un-normalized counts, we observe that the counts in oncogenes are higher in the “cancer-related” datasets compared to the “non cancer-related”. This is particularly true for the ERBB2 gene where counts higher than 1,000 are only present in the cancer-related samples. The tumor suppressor genes have noticeably closer cancer/non-cancer medians abundances, but further investigations such as normalization of counts and improved metadata classification would be needed to draw clear conclusions. Those experiments demonstrate that (even exact) presence/absence queries on such transcripts would not be sufficient to reveal important biological information. The level of expression brings valuable information as shown with the over-expression of oncogenes in cancer-related samples.

## 7 Conclusion

REINDEER is the first and currently only tool that enables scalable indexing of *k*-mer counts in a collection of sequencing datasets. As a byproduct it allows to query the abundances of arbitrary sequences in these datasets. REINDEER relies on the new concept of monotigs, on run-length encoding of count-vectors, optional lower-precision encoding of *k*-mer counts, and on the de-duplication of rows within the count-vectors matrix. To the best of our knowledge and despite their apparent simplicity, neither monotigs nor the greedy construction algorithm described here have previously appeared in the literature. A related idea of breaking unitigs at color changes was used in Kallisto [32]. Note that REINDEER does not encode *k*-mer counts as they exactly appear in a dataset, but instead relies on averaged counts per unitig as an approximation; thus it is not an index that records exact raw counts. REINDEER is able to answer presence/absence queries, although it is not particularly optimized to this task: other techniques such as HowDeSBT, Mantis, and SeqOthello, produce indices that occupy smaller space.

Several exciting future features can be envisioned for REINDEER. We plan on exploring index partitioning (i.e., constructing several REINDEER indices on subsets and merging query results) to accelerate REINDEER’s construction and aim towards indexing SRA-scale collections of datasets. In order to reduce memory usage at the expense of query speed, we are also investigating the possibility of performing on-disk queries.

Dataset quality control is also crucial for some applications such as RNA-seq, in order to draw correct biological conclusions. Removing low abundance *k*-mers before indexing appears to be necessary, as numerous artefactual *k*-mers are created due to sequencing errors. In our experiments with the 2,585 datasets benchmark, we used *k*-mer counts thresholds set in previous literature, but more adaptive thresholds would allow a better compromise between index size and loss of information. Another important aspect is the normalization of returned counts (taking into account dataset size), in order to accurately quantify transcripts, e.g. for the purpose of differential expression analysis.

## Funding

This work was supported by ANR Transipedia (ANR-18-CE45-0020) for CM, DG, MS and RC, by INCEPTION (PIA/ANR-16-CONV-0005) for RC.

## Acknowledgements

Part of the experiments presented in this work were made possible thanks to the EBI computing resources and the Institut Pasteur cluster. The authors would like to thank Antoine Limasset and Maël Kerbiriou for their immense help on BLight software, technical advice. We also thank the whole Transipedia group, and in particular Haoliang Xue and Benoit Guibert for their tests on REINDEER; and finally Paul Medvedev for many helpful comments on the manuscript.

## Appendix

### Statistics on *k* mer size, monotigs and matrix de-deduplication

We used the same 2,512 RNA-seq samples as in the main document to show detailed statistics about REINDEER’s components.

**Table S1:** Comparison of REINDEER’s performances using different *k* values. The datasets used for this experiment correspond to the 2512 datasets that contain sequences of size *≥* 31. 20 threads were used.

| REINDEER | Millions of $k$-mers | | Max Ext. Mem. (TB) | | Wallclock time (h) | | Index size (GB) | | Peak RAM (GB) | |
| --- | --- | --- | --- | --- | --- | --- | --- | --- | --- | --- |
|  | k=21 | k=31 | k=21 | k=31 | k=21 | k=31 | k=21 | k=31 | k=21 | k=31 |
| raw counts | $\sim 3,755$ | $\sim 4,426$ | 6.5 | 4.7 | 39 | 32 | 55 | 47 | 51 | 39 |
| discretized |  |  |  |  | 41 | 30 | 44 | 38 | 43 | 33 |
| $\log_2$ | | | | | 42 | 30 | 32 | 29 | 32 | 31 |
| presence/absence |  |  | 6.3 | 4.5 | 67 | 54 | 29 | 25 | 26 | 25 |

**Table S2:** Details on REINDEER inner components on the 2512 datasets collection, for two sizes of *k* and different count modes. **De-duplication gain** is the ratio of the number of monotigs and of rows of the de-duplicated matrix, showing the gain of having distinct rows instead of one row per monotig. Average monotig size is the average number of *k*-mers in a monotig. Matrices sizes (with/without RLE) are the sizes of matrices stored on disk (16 bits per count), without gzip compression. **RLE gain** shows the gain in matrix size after RLE compression (ratio between size without and with RLE).

|  | Raw counts |  | Log counts |  | Quantized counts |  | Presence/absence |  |
| --- | --- | --- | --- | --- | --- | --- | --- | --- |
|  | k=21 | k=31 | k=21 | k=31 | k=21 | k=31 | k=21 | k=31 |
| Number of distinct $k$-mers (G) | 3.755 | 4.426 | 3.755 | 4.426 | 3.755 | 4.426 | 3.755 | 4.426 |
| Number of monotigs (G) | 1.399 | 1.005 | 1.392 | 0.999 | 1.397 | 1.004 | 1.367 | 0.975 |
| Average monotig size | 2.68 | 4.40 | 2.70 | 4.43 | 2.69 | 4.41 | 2.75 | 4.54 |
| Number of matrix rows (G) | 0.262 | 0.202 | 0.199 | 0.150 | 0.247 | 0.187 | 0.132 | 0.103 |
| **De-duplication gain** | **5.33x** | **4.98x** | **6.87x** | **6.48x** | **5.66x** | **5.36x** | **10.6x** | **9.66x** |
| Matrix size without RLE (TB) | 10.5 | 8.11 | 5.3 | 4.2 | 9.2 | 7.5 | 8.0 | 6.1 |
| Matrix size with RLE (TB) | 0.051 | 0.038 | 0.037 | 0.027 | 0.046 | 0.034 | 0.022 | 0.017 |
| **RLE gain** | **206x** | **213x** | **143x** | **156x** | **215x** | **221x** | **354x** | **353x** |

### Results on metagenomics samples

We tested REINDEER’s performances on another type of data: metagenomic samples^6^. These reads were used as a benchmark experiment in an efficient colored de Bruijn graph publication (VARI [Muggli et al. 2017]) and correspond to cattle metagenomics samples at different time points in a beef production facility [Noyes et al. 2016]. We report REINDEER’s performances on the total 87 datasets, and samples of 10 and 40 datasets. We used REINDEER’s default parameters (*k*=31 and raw counts). On the total dataset, VARI’s authors reported only the peak RAM for their method (246 GB), while REINDEER attains 83 GB. We note that REINDEER’s running time increases following the number of rows in the matrix.

**Table S3:**
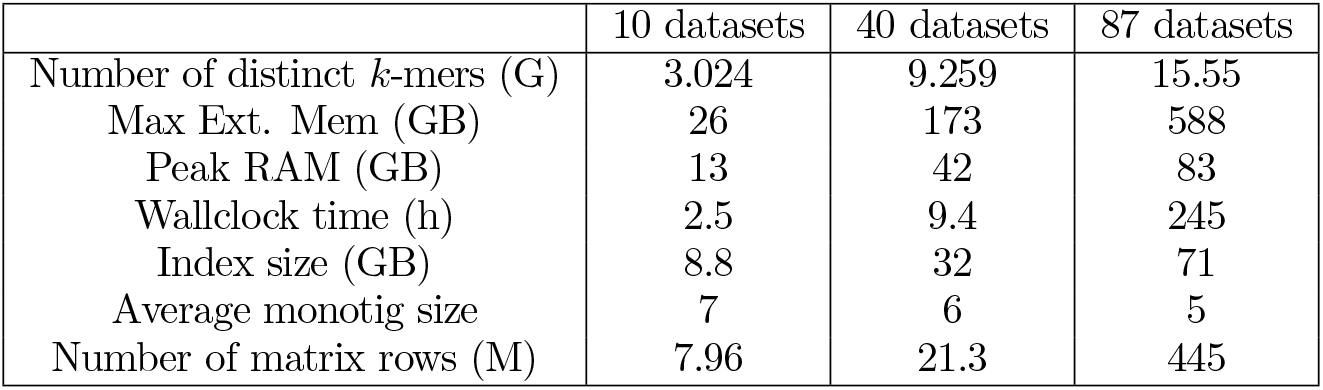
REINDEER on 87 metagenomics samples. Threshold for *k*-mer removal: 2. NA stands for “not accessible”, as this information was not reported in Muggli et al. 2017.

### True *k*-mer counts versus unitig-averaged counts

We selected 15 human RNA-seq datasets^7^ from the SRA by choosing 3 random intervals of 5 datasets from a list of recent Illumina datasets. Figure S1 represents on the *x* axis the average count of the unitig (computed using BCALM2, *k*=31 and minimal *k*-mer abundance of 2), and on the *y* axis the true *k*-mer count (computed using KMC3, *k*=31 and minimal *k*-mer abundance of 2). Each point is thus a pair of numbers (*a,c*) corresponding to a single *k*-mer where the unitig that contains this *k*-mer has an average abundance of *a*, and the true *k*-mer count is *c*. The average Pearson correlation is 0.9985 (standard deviation 0.0026) and average Spearman correlation is 0.9104 (stdev 0.0394) across the 15 datasets.

### Proof of Observation 2

*Proof.* We prove the observation by showing an example (illustrated by Figure 2 second column). Suppose we have two datasets, *d*_1_ and *d*_2_. Dataset *d*_1_ (resp. *d*_2_) contains a single unitig *u*_1_ (resp. *u*_2_). Further assume that *u*_1_ is a prefix of *u*_2_ and *u*_1_ ≠*u*_2_. Observe that *u*_2_ remains a unitig in the union DBG of *d*_1_*∪d*_2_. The first *u*_1_ *k*+1 *k*-mers of *u*_2_ have the same count-vector: the one that contains datasets *s*_1_*,s*_2_ (*s*_1_ and *s*_2_ *>*0). The last |*u*_2_|−|*u*_1_|−*k*+1 *k*-mers of *u*_2_ have count-vector {0*,s*_2_}. It is possible to design sequences for *u*_1_ and *u*_2_ such that the last *k*-mer *x* of *u*_1_ (which is also a *k*-mer in *u*_2_) shares a minimizer with the *k*-mer *y* that follows *x* in *u*_2_. Then the super-*k*-mer that contains both *x* and *y* will be associated to at least two count-vectors: *{s*_1_*,s*_2_*}* through *x* and *{*0*,s*_2_*}* through *y*, hence does not represent a single count-vector.

**Figure S1:**
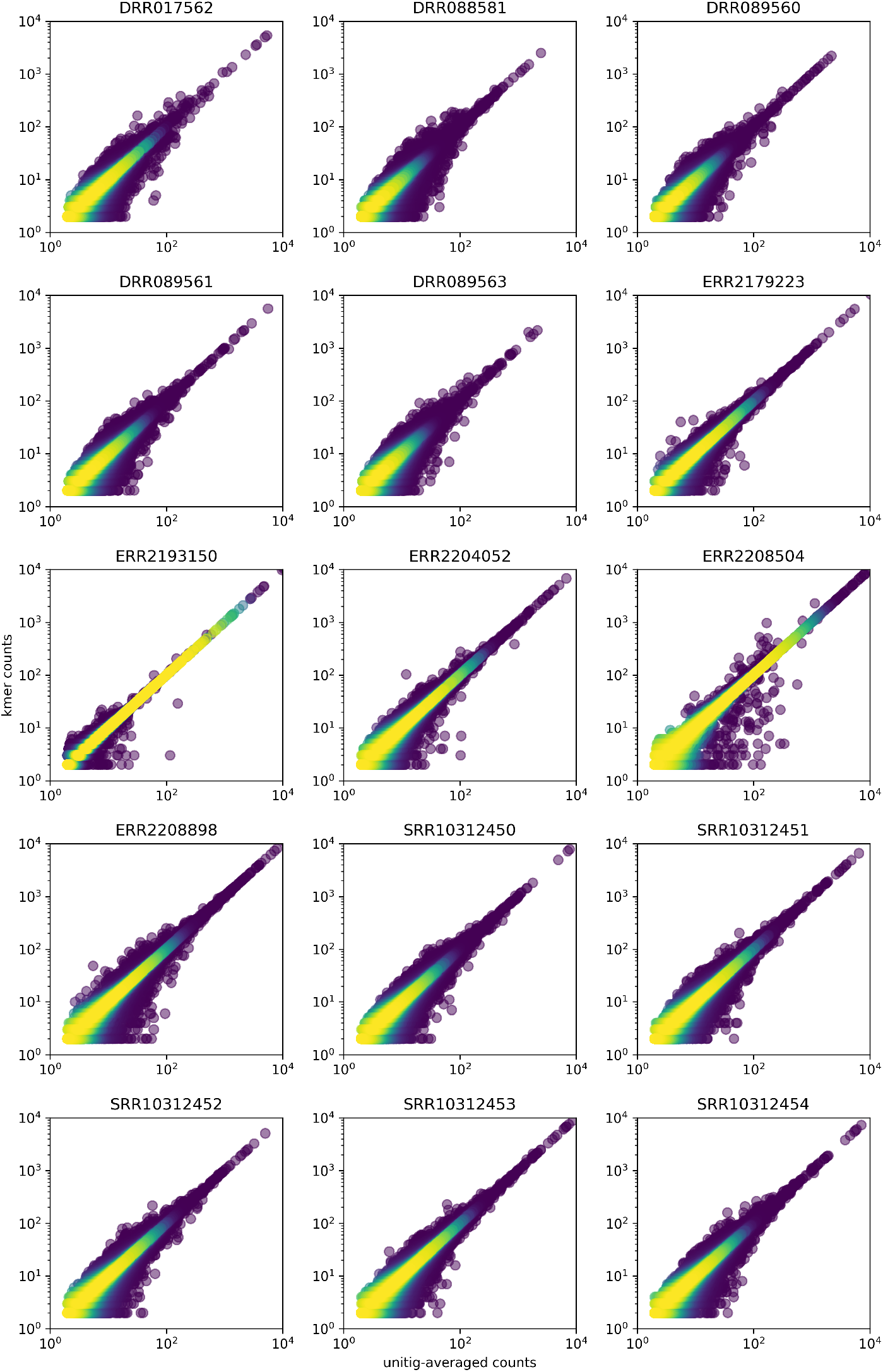
Plots showing correlation between unitig-averaged *k*-mer counts (x) and raw *k*-mer counts (y), for a selection of 15 SRA human RNA-seq datasets. Each plot displays a random subsample of 10,000 *k*-mers per dataset.

**Figure S2:**
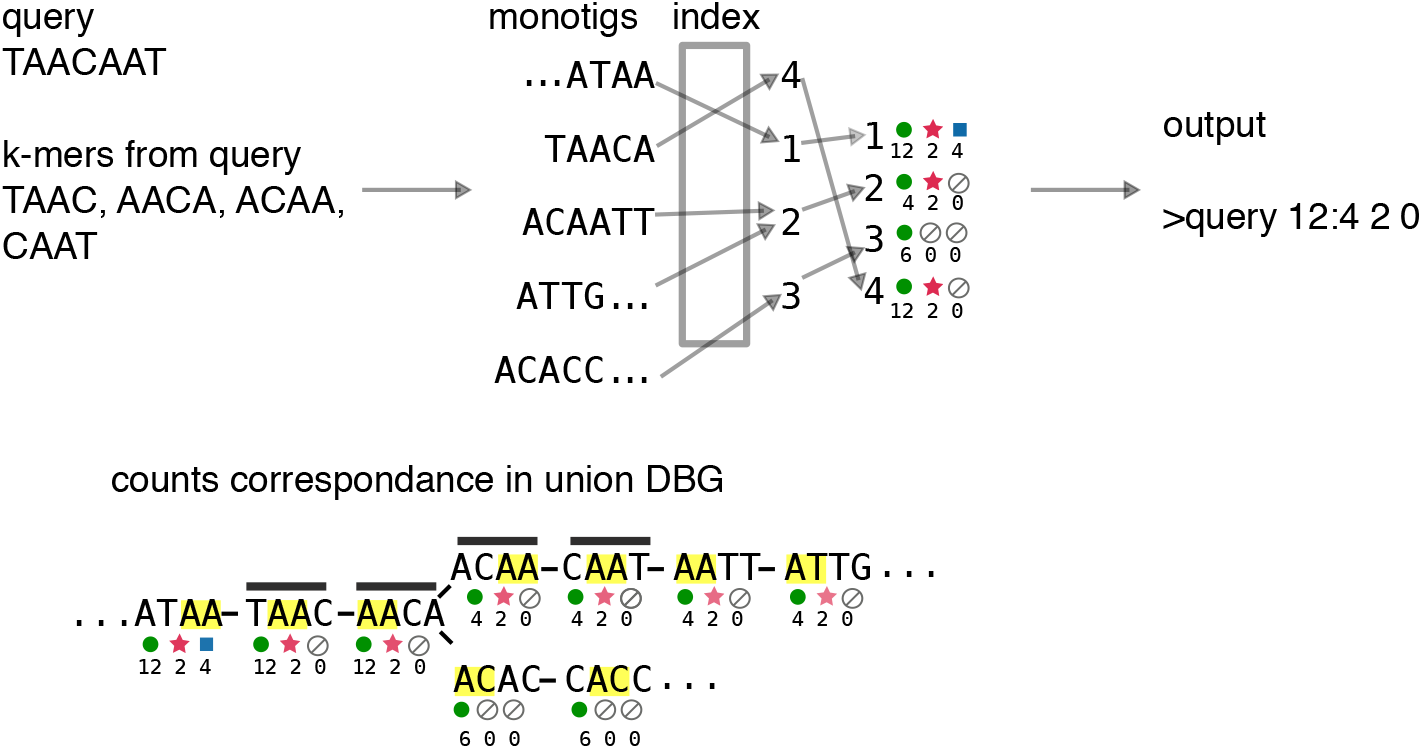
**REINDEER’s output format.** For a query TAACAAT, we start by extracting all distinct *k*-mers. Each *k*-mer is looked for in the index by finding its monotig (TAAC, AACA come from monotig TAACA; ACAA, CAAT come from monotig ACAATT). Monotigs are associated to count vectors. The output shows one row per query, with the query name in the first columns, and then *c* columns for the counts in *c* datasets. The union DBG shows the indexed *k*-mers. Each dataset is represented by a pattern, and counts for this dataset are below the pattern. Since the query sequence can span several unitigs in a given sample graph, different counts can be output for a single dataset. *k*-mers from the query are stressed with dark grey lines. For instance, in the first (round symbol) dataset, the query starts with a count of 12, then switches to 4 because of a bifurcation, therefore we report “12:4” in the output. On the contrary, the counts in datasets 2 (star symbol) and 3 (square symbol) are constant for this query, therefore only one abundance is reported (respectively 2 and 0).

## Footnotes

1 github.com/martinus/robin-hood-hashing

2 Following code by G. Rizk, github.com/GATB/gatb-core/blob/f58d96b/gatb-core/src/gatb/tools/collections/impl/MapMPHF.hpp#L85

3 github.com/powturbo/TurboRLE

4 github.com/kamimrcht/REINDEER/tree/master/reproduce manuscript results

5 the following thresholds, set according to the dataset sizes, were used: https://www.cs.cmu.edu/~ckingsf/software/bloomtree/srr-list.txt

6 https://www.ncbi.nlm.nih.gov/bioproject/292471

7 DRR017562, DRR088581, DRR089560, DRR089561, DRR089563, ERR2179223, ERR2193150, ERR2204052, ERR2208504, ERR2208898, SRR10312450, SRR10312451, SRR10312452, SRR10312453, SRR10312454

## Notes

### Competing Interest Statement

The authors have declared no competing interest.

